# Structure of a human monoclonal antibody in complex with Outer surface protein C (OspC) of the Lyme disease spirochete, *Borreliella burgdorferi*

**DOI:** 10.1101/2024.04.29.591597

**Authors:** Michael J. Rudolph, Yang Chen, Clint Vorauer, David J Vance, Carol Lyn Piazza, Graham G Willsey, Kathleen McCarthy, Beatrice Muriuki, Lisa A. Cavacini, Miklos Guttman, Nicholas J Mantis

**Affiliations:** New York Structural Biology Center, New York, NY; Department of Medicinal Chemistry, University of Washington, Seattle, WA; Division of Infectious Diseases, Wadsworth Center, New York State Department of Health, Albany, NY; Department of Biomedical Sciences, University at Albany, Albany, NY; University of Massachusetts Chan Medical School, Worcester, MA

**Keywords:** antibody, human, epitope, infection, vaccine

## Abstract

Lyme disease is a tick-borne, multisystem infection caused by the spirochete, *Borreliella burgdorferi*. Although antibodies have been implicated in the resolution of Lyme disease, the specific B cell epitopes targeted during human infections remain largely unknown. In this study, we characterized and defined the structural epitope of a patient-derived bactericidal monoclonal IgG (“B11”) against Outer surface protein C (OspC), a homodimeric lipoprotein necessary for *B. burgdorferi* tick-mediated transmission and early-stage colonization of vertebrate hosts. High-resolution epitope mapping was accomplished through hydrogen deuterium exchange-mass spectrometry (HDX-MS) and X-ray crystallography. Structural analysis of B11 Fab-OspC_A_ complexes revealed the B11 Fabs associated in a 1:1 stoichiometry with the lateral faces of OspC_A_ homodimers such that the antibodies are essentially positioned perpendicular to the spirochete’s outer surface. B11’s primary contacts reside within the membrane proximal regions of α-helices 1 and 6 and adjacent loops 5 and 6 in one OspC_A_ monomer. In addition, B11 spans the OspC_A_ dimer interface, engaging opposing α-helix 1’, α-helix 2’, and loop 2-3’ in the second OspC_A_ monomer. The B11-OspC_A_ structure is reminiscent of the recently solved mouse transmission blocking monoclonal IgG B5 in complex with OspC_A_, indicating a mode of engagement with OspC that is conserved across species. In conclusion, we provide the first detailed insight into the interaction between a functional human antibody and an immunodominant Lyme disease antigen long considered an important vaccine target.

## Introduction

Lyme borreliosis or Lyme disease is the most common vector-borne infection in the United States, with an estimated 450,000 cases per year [1]. The primary etiologic agent of Lyme disease is the spirochete bacterium, *Borreliella burgdorferi* sensu latu (herein referred to as simply *B. burgdorferi*). In North America, the spirochete is transmitted to humans by black legged ticks, *Ixodes scapularis* and *Ixodes pacificus*, during the course of a blood meal. The spirochete proliferates at the site of the tick bite, typically resulting in an expanding skin lesion commonly referred to as a bull’s eye rash or erythema migrans [2–4]. In the absence of antibiotic intervention, *B. burgdorferi* disseminates to peripheral tissues, organs, large joints, and the central nervous system, potentially resulting in severe complications including neuroborreliosis, carditis and/or Lyme arthritis [2, 5]. A fraction of Lyme disease patients who receive a full regimen of antibiotics will report persistent health issues (e.g., fatigue, cognitive issues, musculoskeletal pain), a syndrome referred to as post-treatment Lyme disease (PTLD) [6–8].

In an effort to define the immunologic factors that drive resolution of Lyme disease, Blum and colleagues characterized B cell responses in patients across the course of infection [9]. Blood plasmablasts from Lyme disease patients and a healthy cohort were subjected to bulk BCR and single cell paired V_H_ and V_L_ sequencing [9]. The analysis revealed that robust plasmablast responses correlated with a more rapid resolution of disease symptoms. Moreover, expression of a panel of recombinant human monoclonal antibodies (MAbs) using paired V_H_ and V_L_ sequences from representative clones were screened for reactivity with *B. burgdorferi* antigens. The “hits” included a number of immunodominant *B. burgdorferi* surface lipoproteins previously known to be reactive in convalescent human Lyme disease sera, including VlsE, DbpA, DbpB and OspC [9–11]. The recombinant MAbs were further screened for the ability to inhibit growth of *B. burgdorferi* strain B31 in culture. Of the six MAbs with borreliastatic activity (and therefore implicated in disease resolution), two (B2 and B11) were directed against OspC.

Outer surface protein C (OspC) is a member of the small variable surface protein (Vsp) family of immunodominant antigens unique to *Borrelia* and *Borreliella* including the relapsing fever spirochetes (*B. turicatae*) [12]. In *B. burgdorferi*, OspC is expressed during tick transmission and in the early stages of mammalian infection [13–15]. During this period, OspC plays a multifaceted role in pathogenesis, including facilitating spirochete egress from the tick during the course of a blood meal, enabling survival in the early stages of mammalian skin infection, plasminogen interactions, and modulating transmigration across vascular walls [14–20]. The importance of OspC in *B. burgdorferi* pathogenesis is underscored by the fact that active vaccination with recombinant OspC or passive transfer of OspC antibodies completely prevent tick-mediated *B. burgdorferi* infections in mouse models [21–25]. However, while only a single copy of *ospC* is encoded on any *B. burgdorferi* genome, there are >26 *ospC* types and subtypes within *B. burgdorferi* isolates in North America, with multiple types often found in the same geographical proximity [26–28]. The polymorphic nature of OspC is proposed to constitute a means of *B. burgdorferi* immune evasion, as antibodies to one OspC type have limited cross-reactivity and cross-protection with other OspC types [28–31].

Structurally, OspC is ∼21 kDa helical bundle that forms homodimers on the spirochete outer surface [32–36]. Despite a high degree of amino acid variability across the different OspC types, the tertiary and quaternary structures of OspC are virtually identical. As OspC’s N-termini are anchored in the outer membrane via a lipid moiety, the molecule is generally depicted with the stem tethered to the spirochete surface and the dome, with more variable residues, projecting outward [35]. As such it is reasonable to expect that type-specific antibodies target the outward face of OspC, occluding the putative ligand binding cavities [35]. However, the X-ray crystal structure of OspC type A (OspC_A_) complexed with the mouse MAb, B5, reveals a more complicated picture. B5 is the most well-characterized OspC monoclonal antibody originally isolated from *B. burgdorferi*-infected mice [23, 37]. In passive immunization studies, B5 blocks tick-mediated transmission of *B. burgdorferi*, presumably by entrapping the spirochetes in the tick midgut [23]. The X-ray crystal structure revealed that B5 Fabs associates at a nearly perpendicular angle with the lateral sides of homodimeric OspC_A_ [38]. The side-on orientation of B5 with OspC_A_ raises questions about the mechanisms by which the antibody interferes with spirochetal transmission as the putative ligand binding pockets on the outer most face of OspC are unobstructed. A limited insight into the nature of OspC-antibody interactions in mice (and none in humans) represents a gap in our understanding of a critical immune interaction with implications for both Lyme disease resolution and vaccine design. In this study, we report the structure of the human MAb B11 in complex with OspC and its impact on *B. burgdorferi* viability *in vitro*.

## Results

### MAb B11 recognition of native and recombinant OspC

The human MAb, B11, was originally isolated by Blum and colleagues as a recombinant IgG1 with paired V_H_ and V_L_ sequences from single-cell sorted, blood-derived plasmablasts from a Lyme disease patient [9]. The B11 V_H_ and V_L_ sequences are assigned germline HV4-39 and KV3-15, based on the IMGT database[39]. Based on that assignment, the B11 V_H_ and V_L_ coding regions have as many as 16 amino acid mutations from V_H_ HV4-39 germline and 10 from the V_L_ KV3-15 germline, suggesting the antibody has undergone significant somatic hypermutation. Analysis of the available BioProject database indicate that one other B11 clone exists, alongside three closely related clones all derived from the same patient (D. Vance, *unpublished observations*).

The B11 V_H_ and V_L_ coding regions were cloned in-frame into pcDNA3.1-based human IgG_1_ Fc and kappa light chain expression plasmids, respectively, and then co-transfected into Expi293 cells. Purified B11 IgG1 was assessed for the ability to recognize native OspC. Live *B. burgdorferi* strains expressing OspC_A_ (strain B313), OspC_B_ (strain ZS7), or OspC_K_ (strain 297) were incubated with B11 IgG_1_ or an isotype control (PB10 IgG_1_), then subjected to flow cytometry (**Figure 1**). A B31 derivative lacking *ospC* (B31A Δ*ospC*) was also included as a control [40]. By flow cytometry, B11 was highly reactive with strain B313 expressing OspC_A_ (MFI >3300), but not with the *B. burgdorferi* Δ*ospC* mutant. Moreover, MAb B11 did not react with strain ZS7 expressing OspC_B_ (MFI <35) or 297 expressing OspC_K_ (MFI <35), indicating that B11 recognition of OspC is type restricted. Indeed, we employed BLI to examine B11 reactivity with recombinant dimeric OspC_A_. In that assay, B11 IgG had an apparent dissociation constant (K_D_) of ∼39 nM for recombinant dimeric OspC_A_ (**Figure S1**), but negligible reactivity with OspC_B_ or OspC_K_ (**data not shown**). This OspC reactivity profile is reminiscent of MAb B5 [38].

**Figure 1.**
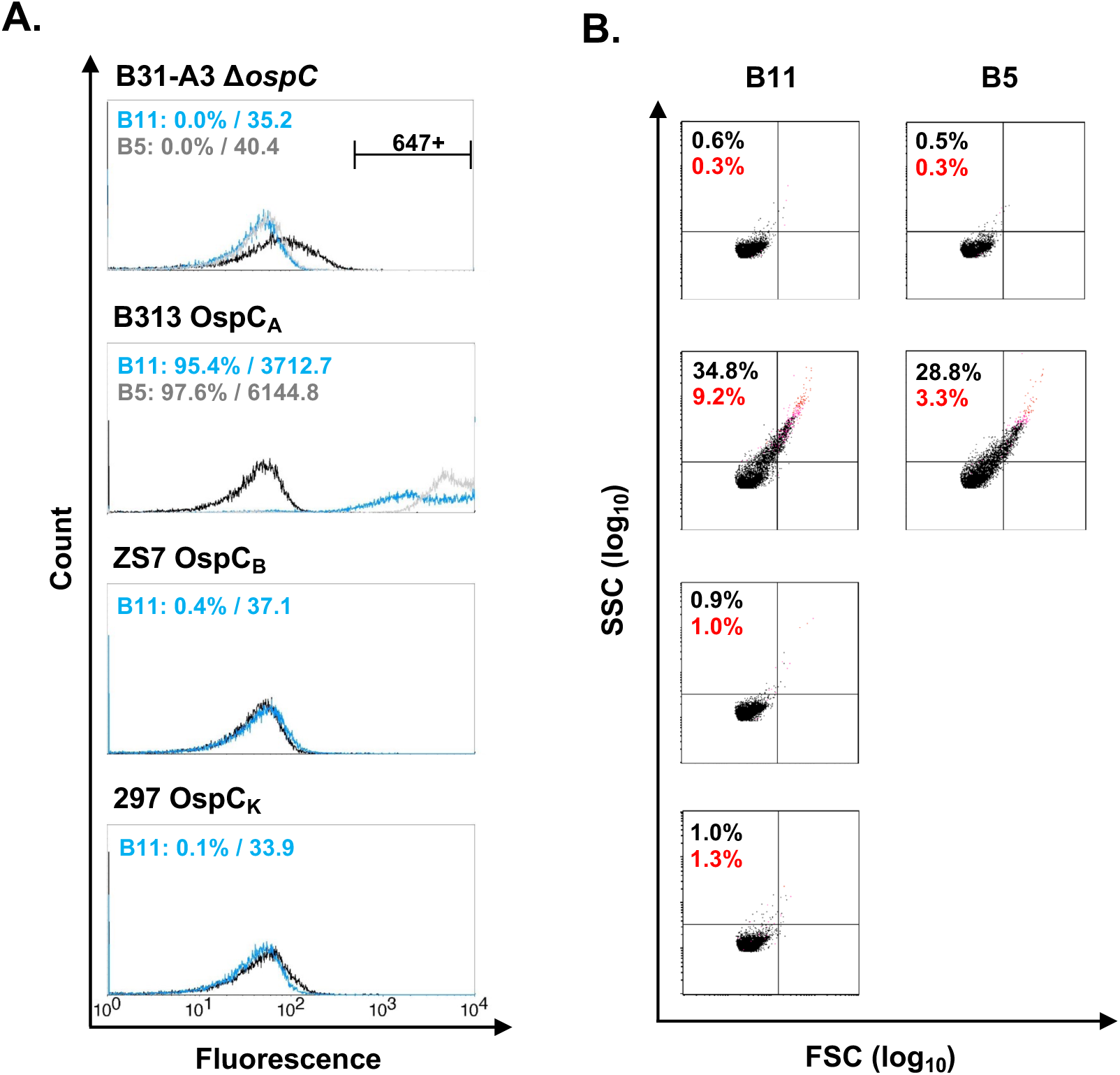
Reactivity of B11 and B5 with OspC_A_ on live spirochetes. Flow cytometric analysis of B11 and B5 MAb reactivity with *B. burgdorferi* strains expressing OspC types A (B313), B (ZS7), or K (297). A B. burgdorferi B31 OspC mutant (B31-A3Δ*ospC*) was included as a control. (A) Representative fluorescence histograms where an Alexa Fluor 647-labeled anti-human IgG secondary antibody was used to detect bacterial surface bound IgG. Panels show B11 (blue) or B5 (gray) compared to the isotype control (PB10, black). Events positive for 647 fluorescence fall to the right of the isotype control. The percent and geometric mean fluorescence intensity (gMFI) of positive events for bacteria incubated with B11 (blue) and B5 (gray) are indicated on each panel. (B) Corresponding FSC/SSC dot plots. Events that are increased in size (FSC) and granularity (SSC) represent multi-bacteria aggregates as the result of antibody-mediated agglutination of live spirochetes. The percentage of events that are agglutinated is indicated (black) and was calculated from the sum of events with increased FSC and SSC, (UL+UR+LR quadrants), relative to the total events counted (20,000). The percent of events positive for propidium iodide staining, indicating membrane permeability, is labeled, and highlighted in red.

### MAb B11 promotes *B. burgdorferi* agglutination and alterations in outer membrane permeability

We recently reported that antibody-mediated agglutination of live *B. burgdorferi* spirochetes results in flow cytometric events with increased size (FSC) and granularity (SSC) [41]. Furthermore, the addition of propidium iodide (PI) to the spirochete preparations just prior to flow cytometry serves as indicator of antibody-induced changes to OM permeability [41]. We have postulated that antibody-mediated agglutination explains, at least in part, how OspA and possibly OspC antibodies compromise *B. burgdorferi* within the tick midgut and limit their transmission to vertebrate hosts [23, 30, 38]. We therefore examined what impact that B11 had on *B. burgdorferi* agglutination. Following B11 treatment, we observed significant *B. burgdorferi* B313 agglutination (20-25% of the population) with a fraction of aggregated cells also PI^+^. This level of agglutination and PI^+^ staining is similar to what we have observed previously for MAb B5 (**Figure 1B**). Neither MAb B11 nor B5 influenced agglutination or PI positivity of the *B. burgdorferi* Δ*ospC* mutant or *B. burgdorferi* strains ZS7 or 297 (**Figure 1B; data not shown**). Thus, B11 recognizes OspC_A_ on the spirochete surface and has the propensity to promote both spirochete agglutination and alterations in outer membrane permeability in the absence of human complement.

### Complement-dependent and independent impact of MAb B11 on *B. burgdorferi* motility

Blum and colleagues reported that MAb B11 (20 μg/mL) had growth inhibitory (bacteriostatic) activity on *B. burgdorferi* in standard culture conditions (BSK II) [9]. To confirm this observation, we examined *B. burgdorferi* B31 motility (as a proxy for cell viability) in the absence and presence of 20% human complement across a range of B11 doses (1-30 μg/mL). To circumvent issues associated with intrinsically low OspC expression by *B. burgdorferi* B31 in culture, we utilized a strain with an IPTG inducible *rpoS* allele, thereby activating native *ospC* expression in *trans* (see Methods). In the absence of complement, B11 had a modest effect on bacterial motility that but did not reach statistical significance as compared to an the isotype control, PB10 (**Figure 2**). The addition of human complement resulted in significant reduction in bacterial motility at B11 concentrations at 3 μg/ml and above. The transmission blocking MAb B5 was significantly more potent than B11 at arresting *B. burgdorferi* motility in absence of human complement and was similar to B11 in the presence of complement (**Figure 2**). Thus, in our hands, B11’s effects on spirochete motility arrest are complement-dependent.

**Figure 2.**
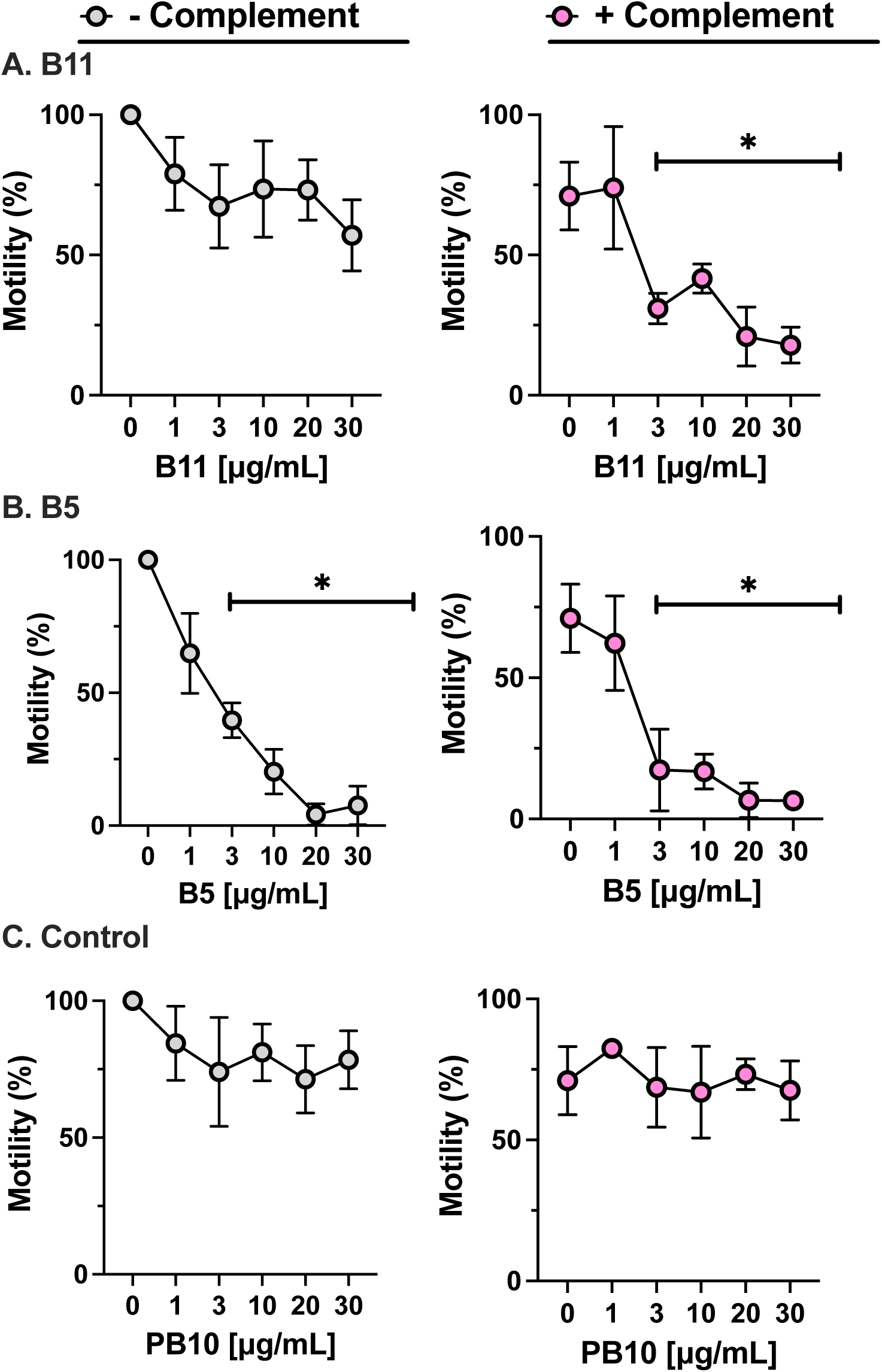
Complement-dependent and -independent effects of B5 and B11. Mid-log-phase cultures of *B. burgdorferi* strain GGW941 treated for 24 h with IPTG induce *rpoS* expression, were adjusted to 5 x 10^6^ bacteria per 50 μL in BSK II without (left columns) and with (right columns) 20% human complement. Cultures were then treated with (**A**) B11, (**B**) B5 or (**C**) PB10 at indicated concentrations (x-axis). Sixteen hours later, the cultures were examined in a double-blind fashion by dark-field microscopy for motile spirochetes. Spirochetes were enumerated in 4 visual fields, and the percent viability was calculated as the ratio of live spirochetes (mean of 4 fields) in treated samples to spirochetes in the untreated control samples (mean of 4 fields). The experimental set up was conducted over the course of three independent sessions and data is plotted as the means for the three days of counting. Statistical analysis to determined significance (*, p<0.05) was done with 2-way ANOVA with Dunnett’s multiple comparisons test for each B11 and B5 versus PB10 (isotype control) at each dose.

### Localization of MAb B11’s epitope on OspC_A_ by HX-MS

We employed HX-MS as a means of identifying B11’s epitope on OspC_A_. A series of preliminary quench and digestion experiments revealed that proteolysis of recombinant dimeric OspC_A_ with Nepenthesin II (without the addition of urea) generated the largest set of observable peptides. Filtering out weak and overlapping signals resulted in 74 unique peptides with a sequence coverage of 98.8% and redundancy of 5.3. The addition of MAb B11 IgG resulted in strong protection at residues 49-57 and 186-196, corresponding to the N- and C-termini of OspC_A_, respectively (**Figure 3; Supplemental Excel file**). Moderate protection was also observed elsewhere (e.g., peptide 157-162). The B11 IgG HX-MS protection profile is reminiscent of the profile we reported for B5 IgG using slightly different HX-MS conditions [38]. To enable a direct comparison between B11 and B5, we subjected OspC_A_ to HX-MS with B5 IgG under the same conditions as B11. The results revealed that HX-MS profiles were indeed distinct with B5 causing notably stronger protection at residues 72-80, 157-176, and 186-196, further from the N/C termini (**Figure 3**). These results suggest that B5 and B11 recognize overlapping but distinct epitopes primarily focused on OspC_A_’s α-helix 5 and 6. To test this experimentally, we performed competitive binding assays between B5 and B11 using the BLI platform in which OspC_A_ was first saturated with B5 then probed with B11. The results confirmed that B11’s ability to associate with OspC_A_ is abrogated by B5 occupancy (**Figure S1C**).

**Figure 3.**
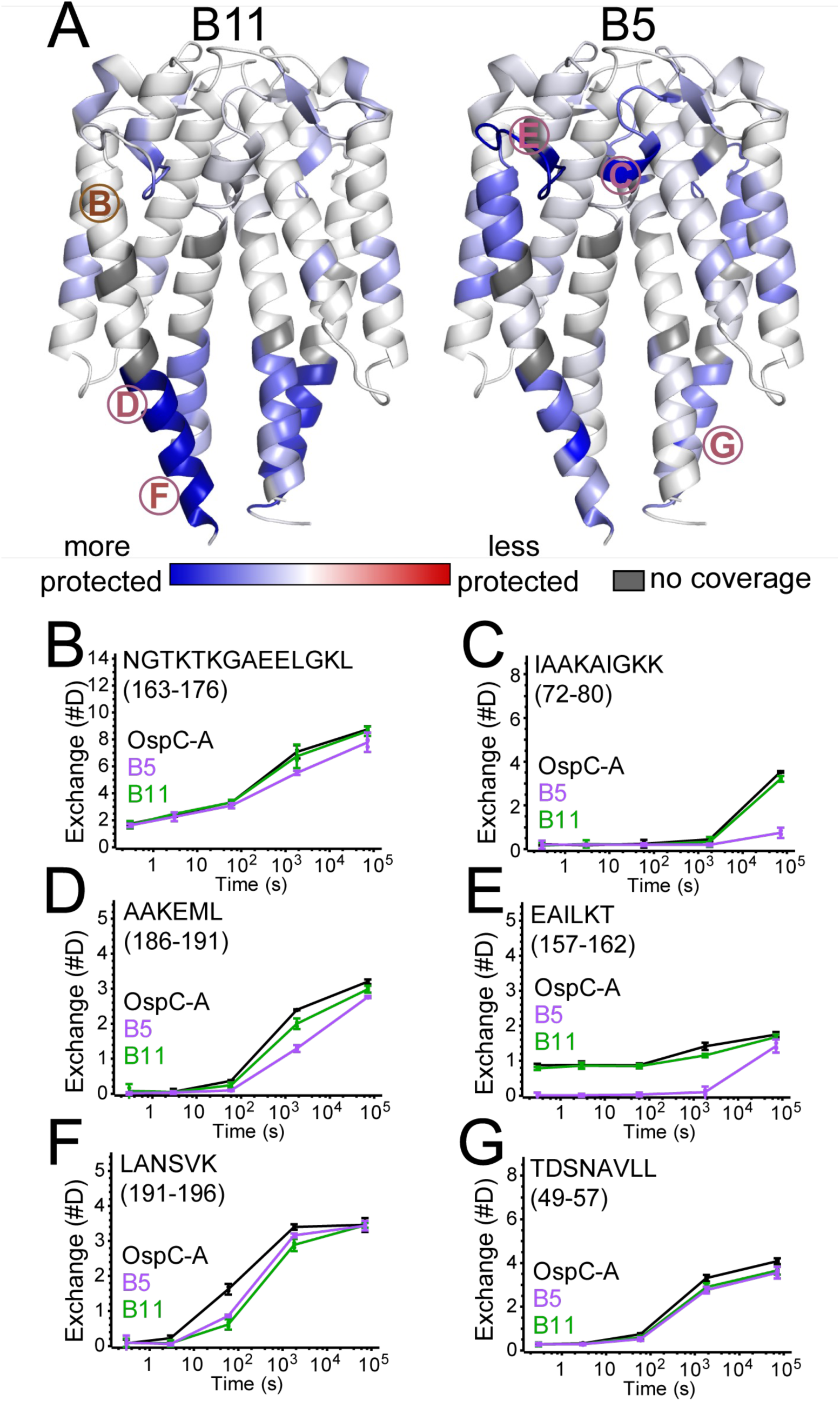
Localization of the B11 epitope on OspC_A_ using HDX-MS. HDX changes upon the addition of (**A**) B11 IgG and B5 IgG are plotted on the structure of OspC_A_ [PDB ID 1GGQ]. Regions with reduced deuterium exchange (more protected) are colored blue, while more deuterium exchange are colored red. (**B-G)** Deuterium uptake plots for unbound OspC_A_ (black liunes), B5-OspC_A_ (purple lines) and B11-OspC_A_ (green lines) are shown for selected regions, with the panel letter corresponding to the labels in panel A. Error bars represent standard deviations from triplicate measurements. Comprehensive exchange data is provided as an Excel file as Supplemental material.

### Structural analysis of the Fab B11-OspC_A_ complex

To resolve B11’s epitope on OspC_A_ in greater detail, we solved the X-ray crystal structure of B11 Fabs in complex with OspC_A_ at 3.1 Å resolution in the P1 space group. The structure revealed two B11 Fabs bound to a single OspC_A_ homodimer (1:1 Fab:OspC_A_ stoichiometry) in a side-on fashion reminiscent (at first glance) of the B5 Fab-OspC_A_ complex (**Figure 4**). The B11 Fab fragments (Fab, Fab’) made identical contacts on opposite sides of the OspC_A_ homodimer (OspC_A-_OspC_A_’), as described in detail below. Each B11 Fab assumed the canonical antibody structure with two heavy chain immunoglobulin domains (V_H_, C_H_1) and two light immunoglobulin domains (V_L_, C_L_) each containing 7-10 β-strands arranged in two β-sheets that fold into a two-layer β-sandwich. The six CDRs (LCDR1-3, HCDR1-3) were situated on one face of the molecule. The four B11 FAb-OspC_A_ complexes within the asymmetric unit were structurally similar, with Root-Mean-Square Deviations (RMSD) in the range of 0.8-1.8 Å upon Cα-superpositioning. OspC_A_ homodimers in the absence and presence of B11 Fabs were also structurally nearly identical [RMSD of 0.6 Å], demonstrating that the B11 Fabs do not induce any significant conformational changes OspC_A_.

**Figure 4.**
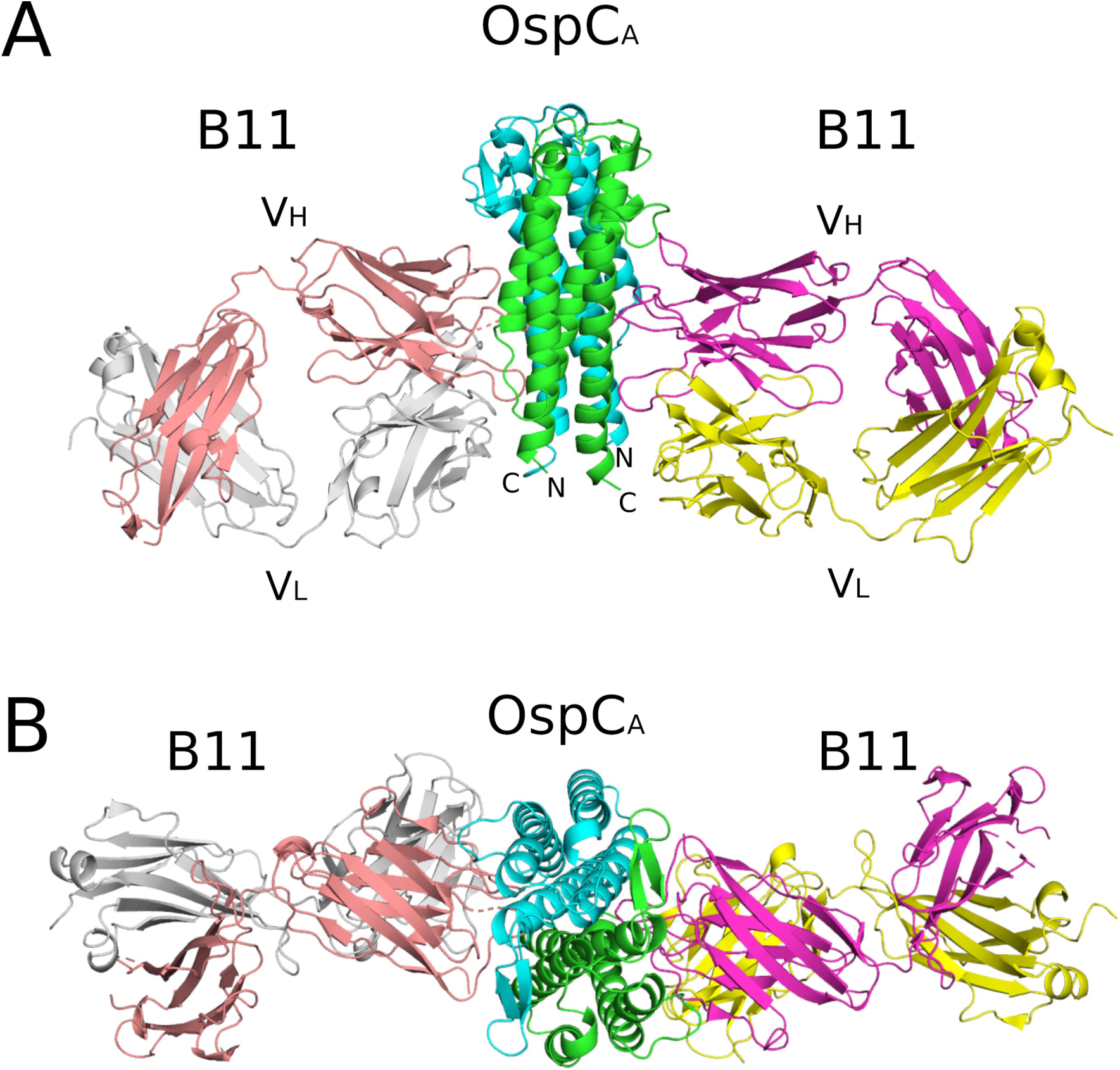
Structure of B11 Fab-OspC_A_. (A) Side-on and (B) top-down ribbon diagrams of OspC_A_ homodimer (OspC_A_, OspC_A_’) in complex with B11 Fabs (B11, B11’). OspC_A_ is colored in cyan and OspC_A_’ in green. The B11 Fab V_H_ and C_H_1 elements are colored in salmon and V_L_ and C_L_ in light gray. The B11’ Fab V_H_ and C_H_1 elements are colored in magenta and V_L_ and C_L_ in yellow. The V_H_ and V_L_ domains (V_H_,V_L_) along with the N- and C-termini of OspC_A_ and OspC_A_’ are labelled accordingly.

Each B11 Fab contacted α-helix 1, α-helix 6, loop 5 (located between α-helices 5 and 6) and loop 6 (after α-helix 6) of their respective OspC_A_ monomers. In addition, the Fabs each bridged the OspC_A_ dimer interface with residues from HCDR1, HCRD3, and LCDR2 engageing with α-helix 1’, α-helix 2’, and loop 2’ on the opposing OspC_A_ molecule (**Figure 5A, B**), thereby demonstrating that B11’s epitope is quaternary in nature. The B11 Fab and OspC_A_ interface buried a total surface area ranging from 2,406 Å^2^ to 2,772 Å^2^ (**Table 2**) Shape complementarity (SC) scores ranged between 0.44-0.59 within the eight Fab-OspC_A_ interfaces in the asymmetric unit, while H-bonds ranged from 3 to 13 and salt bridges from 2 to 4.

**Figure 5.**
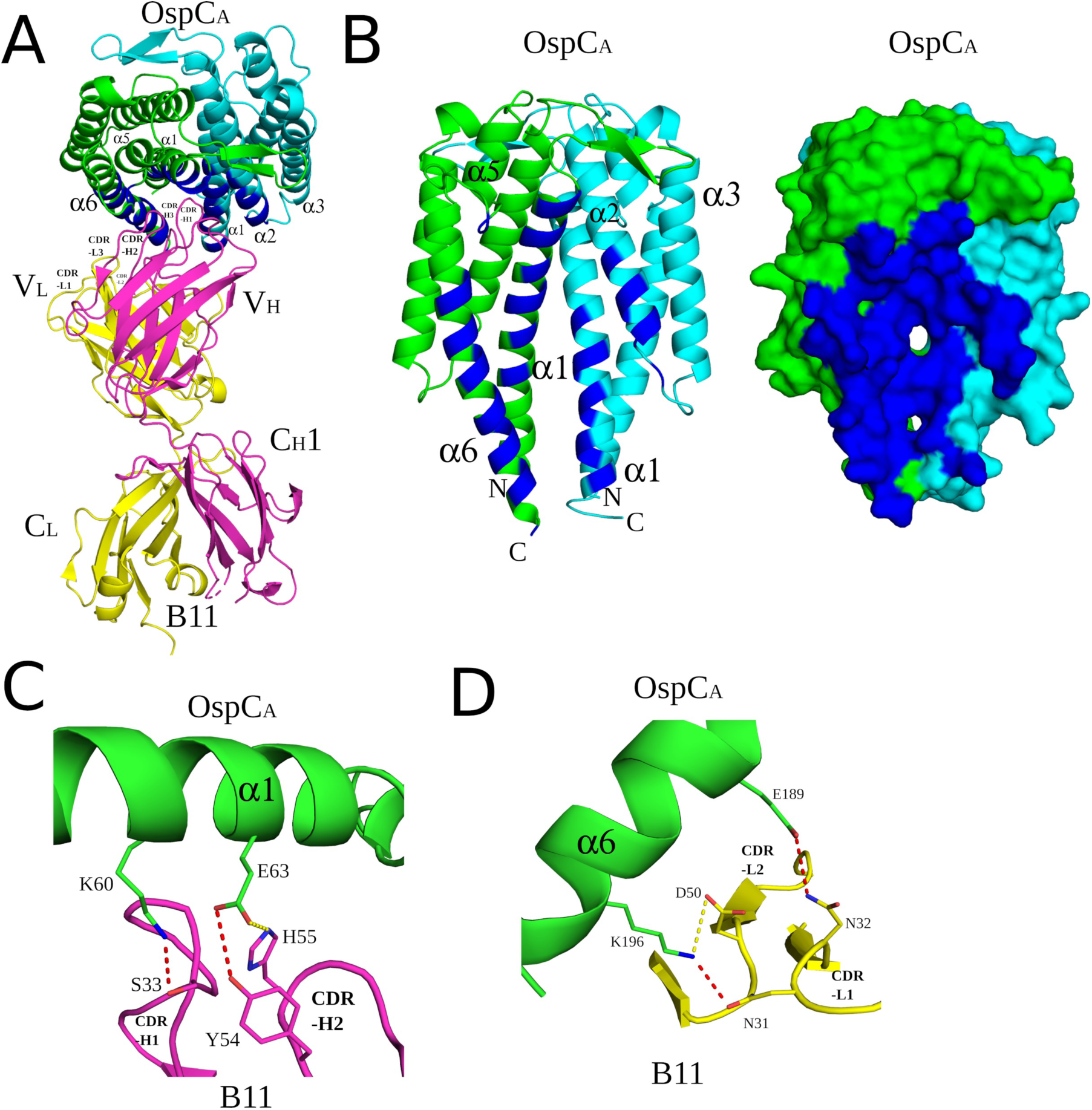
Detailed interactions between B11 and OspC_A_ revealed from the co-crystal structure. (**A**) Ribbon structure (top-down view) of OspC_A_ homodimer (OspC_A_, cyan; OspC_A_’, green) in complex with a single B11 Fab (V_H_ and C_H_1 elements, magenta; V_L_ and C_L_, yellow). The OspC_A_ residues that engage with B11 are colored blue. Key secondary structures (α-helices 1, 2, 3, 5, and 6) and all six CDRs are labeled; (**B**) Ribbon (left) and surface (right) depiction of an OspC_A_ homodimer (OspC_A_, cyan; OspC_A_’, green) with B11-interacting residues shaded in dark blue. OspC_A_ N and C-termini are labelled N and C, respectively. Representations of key H-bonds (red dashes) and salt bridges (yellow dashes) between (**C**) OspC_A_ (green) and B11’ V_H_ domain (magenta) and (**D**) OspC_A_ (green) and B11’ V_L_ domain (yellow). Side chains are drawn as sticks and color coordinated to the main chain color, with nitrogen atoms shaded blue and oxygen atoms shaded red. CDR elements are labelled per convention: CDR-L1, -L3, -L3; CDR-H1, -H2, -H3. OspC_A_ N and C-termini are labelled N and C, respectively. All drawings are depicted as ribbon diagrams except the molecular surface representation in panel B.

**Table 1.** Functional activity associated with MAbs B11 and B5.

|  |  | B11 |  |  |  | B5 |  |  |  |
| --- | --- | --- | --- | --- | --- | --- | --- | --- | --- |
| Strain | type | Binding (%)<sup>a</sup> | Binding (gMFI)<sup>a</sup> | Agg (%)<sup>b</sup> | PI + (%)<sup>c</sup> | Binding (%)<sup>a</sup> | Binding (gMFI)<sup>a</sup> | Agg (%)<sup>b</sup> | PI + (%)<sup>c</sup> |
| *ΔospC* | - | -0.1 +/- 0.1 | -13.8 +/- 10.5 | 0.1 +/- 0.1 | 0.0 +/- 0.2 | 5.5 +/- 12.3 | 55.3 +/- 128.8 | 0.1 +/- 0.2 | 0.2 +/- 0.7 |
| B313 | A | 92.8 +/- 2.0 | 3822.3 +/- 1516.1 | 22.5 +/- 8.6 | 9.3 +/- 5.3 | 94.5 +/- 2.7 | 6903 +/- 1023 | 33.4 +/- 6.1 | 15.2 +/- 10.4 |
| ZS7 | B | 0.3 +/- 0.4 | -7.7 +/- 16.3 | -0.1 +/- 0.1 | -0.1 +/- 0.2 | 0.0 +/- 0.1 | -3.7 +/- 18.1 | -0.1 +/- 0.1 | -0.1 +/- 0.2 |
| 297 | K | 1.5 +/- 3.5 | 0.9 +/- 13.0 | -0.2 +/- 0.3 | 0.01 +/- 0.3 | -0.2 +/- 0.2 | 3.6 +/- 11.1 | -0.1 +/- 0.2 | 0.0 +/- 0.1 |

<sup>a</sup>, *B. burgdorferi* surface binding; <sup>b</sup>, Agglutination; <sup>c</sup>, propidium iodide positive events. Flow cytometric analysis of *OspC* IgG mAbs B11 and B5 and their reactivity with live *B. burgdorferi* strains expressing different *OspC* types. An anti-human IgG Alexa 647-labeled secondary antibody was used to detect mAb binding and is displayed as percentage of events positive for 647 fluorescence, and geometric mean fluorescence intensity (gMFI). The percentage of events with increased size and granularity (FSC/SSC) demonstrating agglutination, and percentage of events with propidium iodide uptake are shown. Displayed are mean values, with background subtraction, +/- standard deviation, n=4.

**Table 2.** Summary of B11-OspC_A_ binding data and interface information.

| **Table 2. Summary of B11-OspC<sub>A</sub> binding data and interface information** |  |  |  |  |
| --- | --- | --- | --- | --- |
| **Interface(s)<sup>a</sup>** | **H-bonds<sup>b</sup>** | **SB<sup>c</sup>** | **SC<sup>d</sup>** | **Total BSA (Å<sup>2</sup>)** |
| 1 (2) | 4 (9) | 3 (3) | 0.51 (0.44) | 2706 (2772) |
| 3 (4) | 12 (13) | 2(3) | 0.48 (53) | 2671 (2710) |
| 5 (6) | 3 (8) | 3(4) | 0.59 (0.47) | 2636 (2814) |
| 7 (8) | 8 (11) | 3(3) | 0.51 (0.59) | 2406 (2600) |
| <sup>a</sup>, primary and secondary (in parentheses) interfaces; <sup>b</sup>,hydrogen bonds; <sup>c</sup>, salt bridges; <sup>d</sup>, shape complementarity score |  |  |  |  |

While all six B11 CDR elements (L1-L3 and H1-H3) associated with OspC_A_, the bulk of the interaction was mediated by HCDR1-3. Specifically, the B11 V_H_ domain accounted for ∼75% of the total BSA within the complex and most of the H-bonds and salt bridges. Key H-bonds include HCDR1 Ser-33 with OspC_A_ Lys-60, and HCDR2 residue Tyr-54 with OspC_A_ Glu-63, with a notable salt bridge occurring between HCDR2 His-55 with OspC_A_ Glu-63 (**Figure 5C**). In terms of the B11 V_L_ domain, H-bonds occurred between LCDR1 Asn-31 and OspC_A_ Lys-196, and LCDR1 Asn-32 with OspC_A_ Glu-189. The B11 V_L_ domain formed a single salt bridge between LCDR2 Asp-50 with OspC_A_ Lys-196 (**Figure 5D**).

### Structural comparisons between B11 and B5 Fab-OspC_A_ complexes

As noted in the Introduction, we recently reported the structure of recombinant homodimeric OspC_A_ complexed with the Fab fragments of the transmission blocking murine MAb, B5 [**PDB ID 7UIJ**] ^26^. The B11 Fab-OspC_A_ complex is reminiscent of that structure in that B5 and B11 Fabs engage with the lateral face of OspC_A_ in a side-on orientation (**Figure 6A**). However, the B5 and B11 Fabs are offset relative to each other, with B5 membrane-distal and B11 membrane-proximal (**Figure 6A**). Moreover, while B5 Fabs are virtually perpendicular to OspC_A_ (**Figure 6A**), B11 Fabs associate with OspC_A_ at a slight angle, thereby positioning B11 Fabs closer to the spirochete outer surface (**Figure 6A**). Nevertheless, there is sufficient distance between B11’s epitope on with OspC_A_ and the spirochete membrane to accommodate an antibody as shown when we super positioned the HIV-1 IgG b12 (PDB ID: **1HZH**) onto the B11 Fab (**Figure 6B**). One possible orientation of OspC-B11 IgG relative to the bacterial outer surface is shown in **Figure S4**.

**Figure 6.**
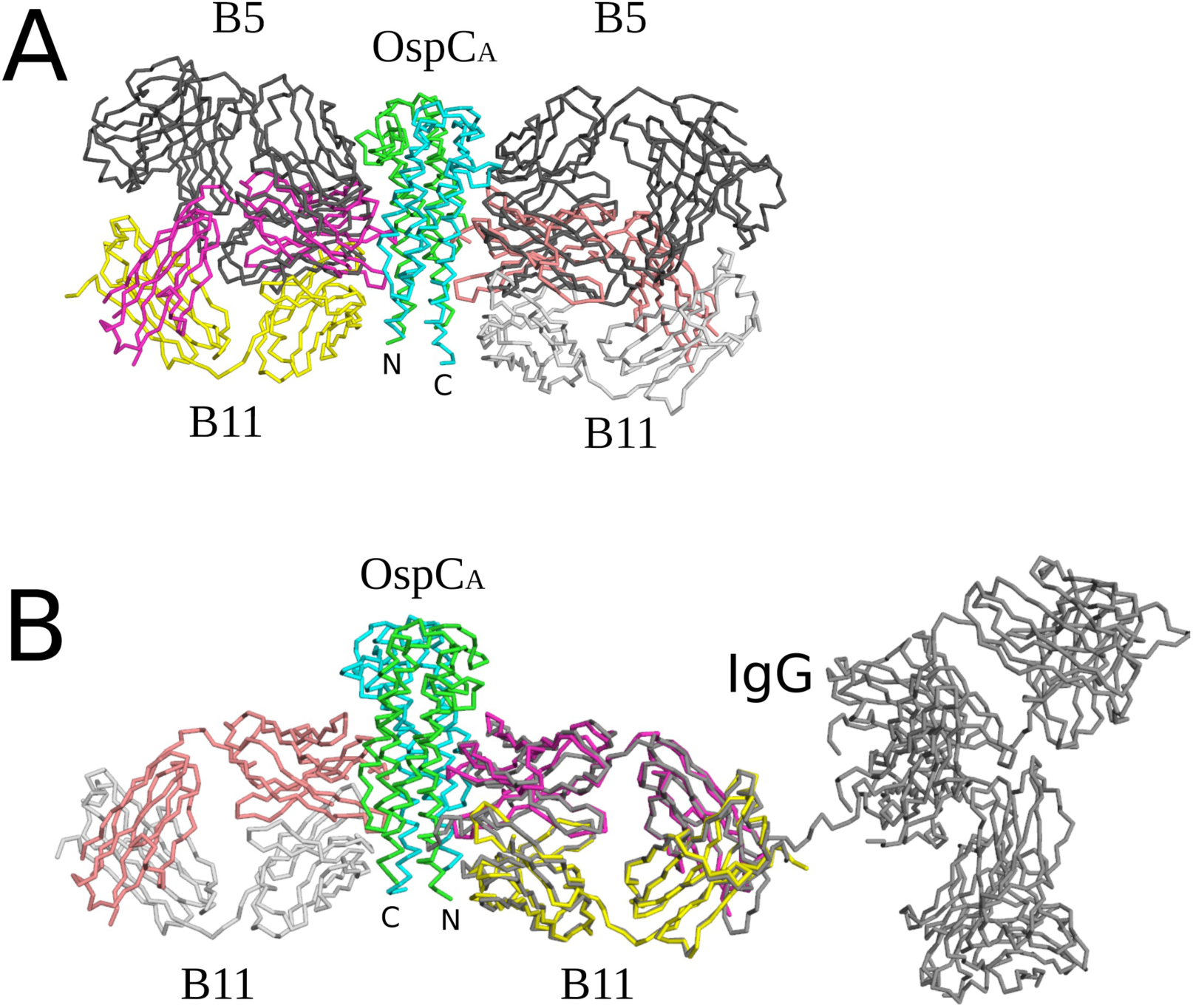
Structural comparison of B11-OspC_A_ with and B5-OspC_A_. (**A**) Superposition of B11-OspC_A_ and B5-OspC_A_ drawn as Cα-traces depicting the similar side-on approach of each Fab to OspC_A_ while highlighting the greater distance of B5 from the N and C-termini of OspC_A_ relative to the B11 position. OspC_A_ is colored cyan and green while the B11 Fab V_H_ and C_H_1 elements are colored in salmon red and V_L_ and C_L_ in light gray. The B11’ Fab V_H_ and C_H_1 elements are colored in magenta and V_L_ and C_L_ in yellow. B5 Fabs are colored dark gray. (**B**) Superposition of the anti-HIV IgG1 mAb (PDB ID 1HZH) drawn as a gray Ca-trace onto B11 also depicted as a Ca-trace revealed no significant steric clash between the OspCA-bound IgG and the spirochete membrane. The OspCA-B11 structure is colored identically to panel A in this figure.

B5 IgG and B11 IgG also resemble each other in terms of OspC type specificity: they react with OspC_A,_ but not OspC_B_ and OspC_K_. To elucidate the structural basis of B11’s specificity for OspC_A_, we superpositioned OspC_A_ from within the B11 Fab-OspC_A_ complex onto the crystal structures of OspC_B_ [**PDB ID 7UJ2**] and OspC_K_ [**PDB ID 7UJ6**], which are structurally similar to OspC_A_ (RMSDs 0.9 Å to 1.1 Å; **Figure S2**). Although the unliganded form of OspC_A_ more closely resembles OspC_K_ (RMSD of ∼0.7 Å) than OspC_B_ (RMSD of 1.0 Å), several structural features of OspC_B_ and OspC_K_ may account for the lack of B11 IgG recognition. The most notable is a Lys at position 113 in OspC_B_ compared to a Gly at residue 114 in OspC_A_. Lys-113 would be expected to sterically clash with B11 Ser-28 (**Figures S3A**, **S4A**). Furthermore, a “deletion” of Ala-74 within α-helix 1 of OspC_B_ relative to OspC_A_ alters the relative configuration of this region. The different conformation of this segment in OspC_B_ would position residue Lys-74 within 4 Å of B11 residue Arg-77 creating an electrostatic clash between these two positively charged residues (**Figures S3B, S5A**). In the case of OspC_K_, the most prominent structural feature damping B11 recognition involves the change of Lys-161 in OspC_A_, which H-bonds to Ser-56 of B11, to Ile-162 in OspC_K_ thus precluding this interaction (**Figures S3C, S5A**).

Understanding the molecular basis of OspC type-immunospecificity is a longstanding challenge with relevance to vaccine design [42, 43]. We previously speculated that B5 IgG reactivity is restricted to just three OspC types (A, C3 and I3), based the variability of key paratope-epitope contacts [38]. By the same token, multiple sequence alignment (MSA) of 23 prominent *B. burgdorferi* OspC types suggest that B11 is similarly restricted. The primary sequences of four of the 23 OspC Types (C3, I3, J, and M) examined contain the residues ostensibly associated with B11 recognition on Type A (**Figure S5B**). OspC types C3, I3, J, and M share 72-80% amino acid identity with OspC_A_. More importantly, sequences within key B11-binding residues within each of these four OspC types are identical OspC_A_. The main-chain structure around residue 74 is conserved (whereas a deletion exists in other types, as described above) and OspC types C3, I3, J, and M contain a glycine or nonbulky amino acid at position 114 alleviating potential steric clash with B11 that appears in other OspC types (as described above). Each of the four selected potential B11-binders possess a phenylalanine at position 177 which buries much of its bulky side chain in the interface with B11. The loss of this bulky side chain would diminish the interaction with B11 by ∼22 Å^2^ based on buried surface area calculations with Phe-177 Å^2^ as a serine residue as found in many of the 23 OspC genotypes. Finally, these four putative B11-binding OspC types also bear a lysine or arginine at residue 161, which putatively H-bonds with B11. From this information, we speculate that B11 likely recognizes four additional OspC types: C3, I3, J, and M (**Figure S5B**). This contrasts with B5 IgG, which we proposed is restricted to OspC types A, C3 and I3 [38]. Cross reactivity with I3 is not surprising given that is a naturally occurring chimera between OspC_F_ and OspC_A_ with the C-terminus (residues 128-199) derived from type A [44].

## Discussion

OspC plays a pivotal role in *B. burgdorferi* tick-to-mammalian transmission and in early stages of dissemination. Although OspC may not be the sole determinant of spirochete dissemination, its importance is underscored by the fact that certain *ospC* genotypes are associated with more invasive *B. burgdorferi* clinical isolates [45–48]. From the host vantage point, *B. burgdorferi* infection is accompanied by a robust OspC-specific antibody response, which subjects the spirochetes to an enormous immune pressure that may contribute to bacterial clearance [42, 49]. Despite the centrality of OspC in both infection and immunity, a high-resolution molecular and structural picture OspC-antibody interactions is lacking in humans.

In this study, we solved the first structure of OspC complexed with Fabs from a human monoclonal antibody. The antibody, B11, was originally isolated by Blum and colleagues as a recombinant IgG1 with paired V_H_ and V_L_ sequences from single-cell sorted, blood-derived plasmablasts from a Lyme disease patient [9]. Recombinant B11 was shown to be bacteriostatic (without addition of human complement) and capable of promoting *B. burgdorferi* opsonophagocytosis in culture [9, 50]. We have extended those observations by confirming that B11 recognizes native OspC on the surface of *B. burgdorferi* strain B31 (but not strains expressing OspC type B or K) and has both complement-independent and -dependent activities against the spirochetes in culture. Based on these observations we predict that “B11-like” antibodies contribute to the reduction and/or clearance of an active *B. burgdorferi* infection in humans.

The crystal structure of B11 Fabs complexed with dimeric OspC_A_ is remarkable in that it demonstrates antibody association with the lateral faces (“stem”) of OspC_A_, rather than the more surface exposed “dome”. Considering that OspC is tethered to the spirochete outer membrane via its lipidated N-terminus, the molecule is generally depicted as projecting perpendicularly from the spirochete surface [33–35]. In such an orientation, the predominantly variable residues face outwards, while conserved amino acids are membrane proximal [34, 35]. Thus, it is natural to assume that type-specific antibodies engage the dome region of OspC via their Fab elements in a top-down orientation [35]. That assumption was called into question by the structure of B5 Fab-OspC_A_ [38], and further undermined by the structure of B11 Fab-OspC_A_.

Perhaps it is worth turning things on their side when rethinking the nature of OspC-antibody interactions. Indeed, Lawson and colleagues pondered this exact possibility when scrutinizing the structures of OspC and related Vsps from *B. turicatae* [35]. They cite evidence to suggest that the N- and C-termini of OspC and Vsp1 dimers “…either adopt a mostly ‘random coil’ conformation or are disordered.” If correct, they argue, then OspC is essentially bound to the spirochete by a flexible tail, thereby enabling the bulk of OspC to sit sideways or even “upside down” on the cell membrane. Moreover, they note that such an unconventional orientation would explain how IgM antibodies are able access the very C-terminal decapeptide of OspC, a linear epitope known as C10 [51]. In such a configuration, the Fc regions of antibodies like B11 IgG would protrude from the bacterial surface and be prime targets for components of the complement system and Fc receptors on phagocytes [50].

The structural elucidation of the of B11 Fab-OspC complex has implications when considering the design of OspC-based vaccine antigens. First, the B11 Fabs bridge the OspC-OspC’ dimer interface. This observation indicates that B11 underwent affinity maturation in the context of dimeric (not monomeric) OspC. From the standpoint of vaccine design, preserving the dimeric interface of OspC antigens is paramount for eliciting potent B11- and B5-like antibodies, an observation that has been alluded to by others [52]. Second, a detailed view of B11 and B5’s epitopes reveal the molecular basis of OspC type-specificity, a factor that has limited OspC’s utility as a Lyme disease vaccine antigen [28, 42, 53]. However, considering the current exponential advances in protein modeling and design, the availability of structural B cell epitopes will prove invaluable. Finally, tt is worth noting that in separate efforts we have identified human monoclonal and camelid-derived single-domain antibodies that have pan-OspC reactivity (L. Cavacini, D. Vance, M. Rudolph, N. Mantis, *manuscripts in preparation*). High-resolution epitope mapping studies of those pan-reactive OspC antibodies are ongoing, alongside passive protection studies in mouse models of *B. burgdoferi* by tick-mediated infection.

## Materials and Methods

### Cloning and expression of B11 IgG

The B11 V_H_ and V_L_ coding regions were extracted from NCBI’s BioProject database under accession PRJNA470931 (GSE114310) [9]. The B11 V_H_ and V_L_ coding regions were synthesized as gBlock™ dsDNA fragments (GenScript, Piscataway, NJ) and cloned in-frame into pcDNA3.1-based human IgG_1_ Fc and kappa light chain expression plasmids, respectively [54, 55]. The two plasmids were co-transfected into Expi293 cells using ExpiFectamine293™ transfection reagents (Thermo Fisher Scientific, Waltham, MA), following manufacturer’s instructions. Five to six days later, supernatants containing the secreted antibodies were harvested and clarified before being subjected to protein A chromatography. Affinity purified IgG_1_ B11 was then subjected to buffer exchange into PBS and stored at 4 °C. B11 Fabs were generated as described [38]. Purification and specificity of MAbs B5 [38]and hPB10 [56]have been previously described.

### Cloning, expression, and purification of OspC

Recombinant OspC_A_ (residues 38 to 201) from *B. burgdorferi* B31 [UniProt: accession Q07337] was expressed and purified as a homodimer from *E. coli*, as described [38].

### *B. burgdorferi* strains and culture conditions

*B. burgdorferi* strains expressing OspC types A (B313), B (ZS7), K (297), and the *ospC* deletion strain B31A3ΔospC were cultured in BSK-II medium at 37°C with 5% CO_2_ to mid-log phase [57], collected by centrifugation (3,300 × *g*), washed, suspended in BSK II with 20% glycerol and stored at −20 °C until needed. Strains B313 [40], ZS7, and B31A3Δ*ospC* (“*ospC*K1”) were kindly provided by Dr. Yi-Pin Lin (Wadsworth Center). Strain 297 was obtained from the American Type Culture Collection (ATCC; Manasas,VA).

A strain of *B. burgdorferi* B31 was engineered to deliver consistent ectopic expression of *ospC* under *in vitro* culturing conditions (BSK-II at 37℃). An IPTG-inducible *rpoS* expression plasmid was created through modification of the pBSV2G-derived *mCherry* plasmid, pGW163 [58]. Briefly, the *rpoS* open reading frame (ORF) was amplified from *B. burgdorferi* B31 [NCBI:txid224326] genomic DNA using Q5 DNA polymerase (NEB, Beverly, MA) and the b31_rpoS_gibson_F (5’-agaattcattaaagaggagaaattacccatgaacatatttagtaatgaggatttaaacat −3’) and b31_rpoS_gibson_R (5’-gtaaaacgacggccagtgccaagcttaatttatttcttcttttaattttttaagaactc-3’) tailed primer set. The *mCherry* ORF was excised from pGW163 using NcoI and HindIII (NEB) and subsequently replaced with the tailed *rpoS* DNA fragment using the HiFi DNA assembly kit (NEB). Following the assembly reaction, the DNA was transformed into NEB^®^ 5-alpha F’ I q chemically competent cells (NEB). Recombinants were selected on LB agar supplemented with 10 µg/ml of gentamicin. Transformants that arose following overnight incubation at 30℃ were screened via PCR using the b31_rpoS_gibson_F and b31_rpoS_gibson_R primer set. The IPTG-inducible *rpoS* expression plasmid, pGW181, was isolated from a single PCR-positive colony. Following purification from a large batch culture, pGW181 was electrotransformed into *B. burgdorferi* B31 as described [59], ultimately yielding the IPTG-inducible *rpoS* expression strain, GGW941.

For routine culture, frozen aliquots of *B. burgdorferi* were thawed and used to inoculate modified BSK-II lacking gelatin, supplemented with 6% rabbit serum (Pel-Freeze Biologicals, Rogers, AR) [60]. Base BSK-II medium was prepared by the Wadsworth Center’s Tissue and Media Core Facility and filter sterilized (0.2 μm) prior to use. Cultures were maintained at 37°C with 5% CO_2_ and passaged by dilution (1:10,000) into fresh BSK-II medium. *B. burgdorferi* cultures were routinely inspected for culture viability and motility during *in vitro* culture maintenance prior to the initiation of any experiments.

To induce RpoS expression, GGW941 culture medium was spiked with 250 µM IPTG at 24 h before use. For flow cytometry and microscopy studies (as described below), *B. burgdorferi* cells were collected in mid-logarithmic phase (∼1-3 × 10^7^ spirochetes per mL). As controls, spirochetes were heat killed by incubation at 56°C for at least 30 min, as described [61].

### *B. burgdorferi* motility determinations by dark field microscopy

Mid-log-phase cultures of GGW941 treated 24 h prior with IPTG to induce *rpoS* expression, were adjusted to 5 x 10^6^ bacteria per 50 μL in PBS then treated with indicated OspC MAbs (1, 2, 3, 10, 20, 30 μg/mL) in the presence or absence of 20% human complement (Sigma-Aldrich) for 16 or 24 h. Cultures were examined in a double-blind fashion by dark-field microscopy for motile spirochetes. 1:5 dilutions of the reactions were prepared by aspirating from the center of the reaction and diluted into fresh BSK-II medium. Dark-field microscopy was performed utilizing a Trinocular DF microscope (AmScope) equipped with a camera with reduction lens (AmScope SKU: MU1603) using a 40× dry darkfield condenser (AmScope; DK-DRY200). Spirochetes were considered dead when complete loss of motility and refractivity was observed. Spirochetes were enumerated in 4 visual fields, and the percent viability was calculated as the ratio of live spirochetes (mean of 4 fields) in treated samples to spirochetes in the untreated control samples (mean of 4 fields). Polyclonal serum from *B. burgdorferi*-infected mice and MAb B5 were used as positive controls; naive serum and the PB10 isotype were used as negative controls. This experimental set up was conducted over the course of three independent sessions and data is plotted as the means for the three days of counting. Statistical analysis was determined using 2-way ANOVA with Dunnett’s multiple comparisons test in which experimental means (B11, B5) were compared to control mean (PB10) for each antibody concentration.

### Flow Cytometry

To examine the ability of B11 and B5 IgG1 to bind the spirochete surface and promote bacterial agglutination, and membrane permeability, *B. burgdorferi* strains expressing OspC types A (B313), B (ZS7), and K (297), were cultured, treated with 10 µg/ml IgG1, and analyzed by flow cytometry as described [38]. Briefly, an Alexa Fluor 647-labeled goat anti-human IgG [H+L] secondary antibody (Invitrogen) was used to detect bound IgG1 to the bacterial surface, and propidium iodide staining was used to indicate bacterial membrane permeability. The *ospC* deletion strain B31-A3*ospC*K1was used as a negative strain control, and the ricin toxin MAb, PB10, was used as an IgG1 isotype control. Bacteria were analyzed on a BD FACSCalibur (BD Biosciences). Voltage was set with untreated bacteria as a reference point (gMFI∼ 5.0), and bacteria were gated on forward scatter (FSC) and side scatter (SSC) to exclude debris and assess aggregate size and granularity. Agglutination was calculated as the sum of events in the upper-left, upper-right, and lower-right quadrants relative to the total event counted (20,000).

### Epitope mapping of B11 by HX-MS

Stock concentrations of OspC_A_ (8.5 µM) in PBS alone or in a complex with a 2-fold molar excess of antibody B11 were diluted into 90 µL of deuterated PBS buffer (20 mM phosphate, 150 mM NaCl, 0.02% sodium azide, 1 mM EDTA pH* 7.54, 85%D final) containing 0.2 nM bradykinin and incubated 3 sec on ice, or either 3 seconds, 1 minute, 30 minutes, or 20 hours at 21°C. Each starting stock also included a mixture of imidazolium compounds to serve as exchange reference standards [62]. At the desired time point the sample was rapidly mixed with an equal volume of ice cold 0.2% formic acid and 0.1% trifluoroacetic acid (TFA) for a final pH of 2.5. Samples were then immediately frozen on ethanol/dry ice and stored at −80°C until LC-MS analysis. Undeuterated samples were prepared the same way but with undeuterated buffer for each step.

Samples were thawed at 5°C for 8 minutes and injected using a custom LEAP robot integrated with an LC-MS system [63]. The protein was first passed over a Nepenthesin II column (2.1 x 30 mm; AffiPro) at 400 µL/min for inline digestion with the protease column held at 20°C. Peptides were then trapped on a Waters XSelect CSH C18 trap cartridge column (2.1 x 5 mm 2.5 µm) and resolved over a CSH C18 column (1 x 50 mm 1.7 µm 130Å) using linear gradient of 5 to 35% B (A: 0.1% FA, 0.025% TFA, 5% ACN; B: ACN with 0.1% FA) over 10 minutes and analyzed on a Thermo Orbitrap Ascend mass spectrometer at a resolution setting of 120,000. A series of washes over the trap and pepsin columns was used between injections to minimize carry-over as described [63]. Data dependent MS/MS acquisition was performed on an undeuterated sample using rapid CID and HCD scans and processed in Byonic (Protein Metrics) with a score cutoff of 150 to identify peptides. Deuterium incorporation was analyzed using HDExaminer v3 (Sierra Analytics).

### Crystallization and data collection

Fab B11 was complexed with dimeric OspC_A_ in a 1:1 stoichiometry, then concentrated to 10 mg/ml final for all crystallization trials. Crystals were grown by sitting drop vapor diffusion using a protein to reservoir volume ratio of 1:1 with total drop volumes of 0.2 μl. Crystals of the B11 Fab-OspC_A_ complex were produced at 22°C using a crystallization solution containing 100 mM sodium HEPES [pH 6.5], 18.6% PEG 6K, and 6.6 mM prasseodymium acetate. All crystals were flash frozen in liquid nitrogen after a short soak in the appropriate crystallization buffers supplemented with 10% ethylene glycol. Data were collected at the 24-ID-E beamline at the Advanced Photon Source, Argonne National Labs. All data was indexed, merged, and scaled using HKL2000[64] then converted to structure factor amplitudes using CCP4 [65].

### Structure determination and refinement

The B11 Fab-OspC_A_ complex structure was solved by molecular replacement using Phaser [64]. Molecular replacement calculations were performed using the V_L_ and C_L_ domain coordinates of the human monoclonal mAb MAB4-5 (PDB ID: 5Y11) along with the V_H_ and C_H_1 domains from the human anti-HIV antibody (PDB ID: 5UBY) as the search model for the first two B11 Fabs in B11-OspC_A_ complex within the asymmetric unit. The OspC_A_ coordinates (PDB ID: 1GGQ) were then used as the search model for the OspC_A_ dimer in the first OspC_A_-B11 complex. The initial B11-OspC_A_ complex detected was then used to find the additional three B11-OspC_A_ complexes present in the asymmetric unit. The resulting phase information from molecular replacement was used for some manual model building of the B11-OspC_A_ structure using the graphics program COOT [66] and structural refinement employing the PHENIX package [67]. Data collection and refinement statistics are listed in **Table S1**, as is the Protein Data Bank (http://www.rcsb.org/pdb/) code for the B11-OspC_A_ structure PDB ID 9BIF. The PDF validation report is provided in Supplemental Information. Molecular graphics were prepared using PyMOL (Schrodinger, DeLano Scientific LLC, Palo Alto, CA).

## Data availability

All results and data files associated with this article are included in the main text and supplemental material.

## Supporting information

This article contains supporting information, including supplemental tables with X-ray crystallography conditions, supplemental figures related to epitope mapping, and HX-MS data files and PDB validation report.

## Acknowledgements

We are grateful to Drs. Lisa Blum and William Robinson (Stanford University) for sharing the B11 VH and VL sequences and for follow-up discussions. We thank Dr. Renjie Song at the Wadsworth Center’s Immunology Core facility for assistance with flow cytometry. We thank the DNA sequencing core and members of the Wadsworth Center’s bioinformatics core for useful discussions. As always, we are grateful to Elizabeth Cavosie (Wadsworth Center) and Donna DeLuca (Health Research, Inc) for administrative assistance and program overview.

## Funding sources

This work was supported by the National Institute of Allergy and Infectious Diseases (NIAID), National Institutes of Health (NIH), Department of Health and Human Services Contract No. 75N93019C00040. X-ray analysis as conducted at the Northeastern Collaborative Access Team beamlines, which are funded by P30 GM124165 from the National Institute of General Medical Sciences (NIGMS), NIH. The Eiger 16M detector on the 24-ID-E beam line is funded by a NIH-ORIPHEI grant (S10OD021527). HDX-MS studies conducted at the University of Washington was supported by award S10OD030237 from NIGMS. The content is solely the responsibility of the authors and does not necessarily represent the official views of the NIH.

## Conflict of interest

The authors have no conflicts of interest to declare.

## Notes

### Competing Interest Statement

The authors have declared no competing interest.

### Summary of Updates

Minor edits and corrections have been made to Results and Figures. The final PDB validation report is also now available.

