## Supplemental Tables and Figures for "Structure of a human monoclonal antibody in complex with Outer surface protein C (OspC) of the Lyme disease spirochete, *Borreliella burgdorferi*"

### Table S1

| **Table S1. Data collection and refinement associated with OspC$_A$-B11 Fabs** | |
|---|---|
| **Data Collection** | |
| Complex | OspC$_A$-B11 |
| Space group | P1 |
| Cell parameters: *a,b,c* (Å) / α β γ (°) | 96.7, 97.3, 168.3 / 97.6, 90.3, 106.2 |
| BNL Beamline | 21-ID-E |
| Resolution range [a,] (Å) | 50-3.10 (3.15-3.10) |
| wavelength (Å) | 0.979 |
| No. of reflections | 3686979 |
| Average redundancy[a] | 3.1 (3.2) |
| $(I)/(\delta)$[a] | 7.6 (1.6) |
| Completeness[a] (%) | 90.8 (90.6) |
| $R_{merge}$[a,b] (%) | 20.0 (83.3) |
| $CC_{1/2}$[a,c] | (0.51) |
| **Refinement** | |
| Bragg spacings [a,] (Å) | 48.6-3.1 (3.13-3.09) |
| $R$[d] / $R_{free}$[e] (%) | 23.5 / 29.4 |
| No. of Protein atoms | 35,323 |
| No. of Waters | 193 |
| RMSD bond length (Å) | 0.005 |
| RMSD bond angle (°) | 1.08 |
| Ramachandran favored / allowed[f] (%) | 98.6/ 99.6 |
| PDB code | 9BIF |

[a] Values in outermost shell are given in parentheses.
[b] $R_{merge} = (\sum | I_i - \langle I_i \rangle |)/\sum | I_i |$, where $I_i$ is the integrated intensity of a given reflection.
[c] $CC_{1/2} = (1+q^2\sigma_\varepsilon^2/\langle I \rangle 2^{-1}$ , where $\sigma_\varepsilon$ denotes the mean error within a half-datase, $CC_{1/2}$ is the correlation coefficient of two split data sets each derived by averaging half of the observations for a given reflection.
[d] $R = \sum | |F_o| - |F_c| | / \sum | |F_o| |$, where $F_o$ and $F_c$ denote observe and calculated structure factors, respectively.
[e] $R_{free}$ was calculated using 5% of data excluded from refinement.
[f] Calculated using Molprobity.

### Figure S1

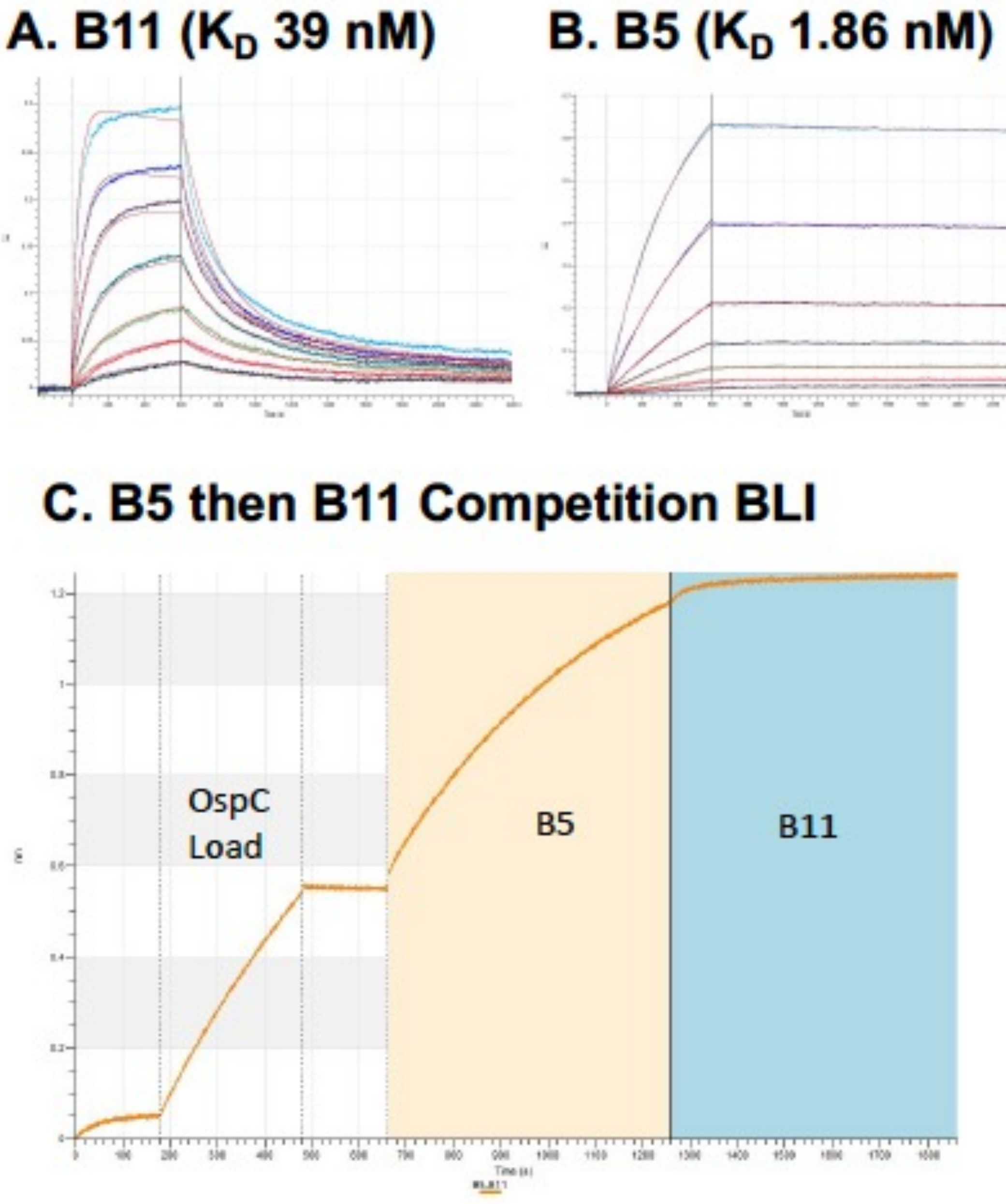


**Figure S1. B5 and B11 binding affinities and competive binding by BLI.** (A) Biolayer interferometry sensorgram of B11 affinity for OspC$_A$. Concentrations range from 100 nM B11 down to 1.56 nM. (B) BLI sensorgrams of B5 affinity for OspC$_A$, at the same concentrations as B11. (C) Competition BLI experiment in which B5 (yellow) at 100 nM was allowed to bind to OspC$_A$ loaded on the sensors, and then subsequently exposed to B11 (blue). Due to B11's >10-fold higher affinity for OspC$_A$ compared to B11 (1.86 nM versus 39 nM), the OspC sensors were first immersed in B5 solution. As predicted, the results demonstrated that occupancy of OspC$_A$ with B5 inhibited subsequent binding by B11, based on the lack of a secondary (RU) when immersed in B11 solution.

### Figure S2

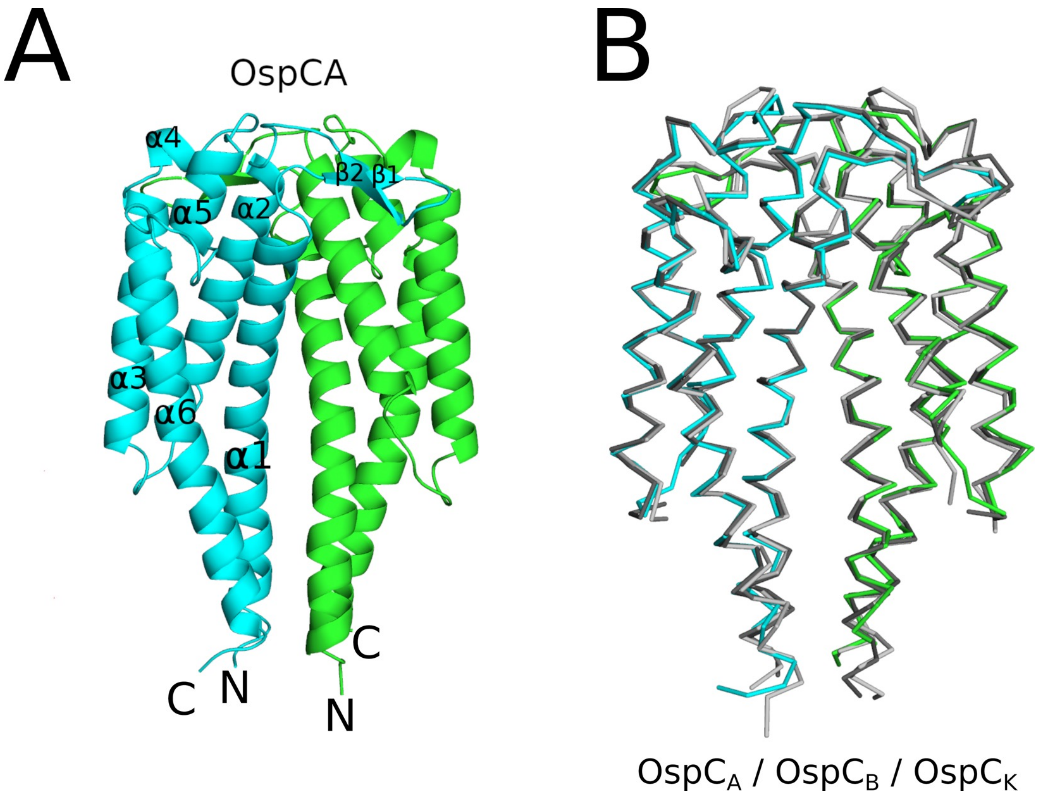


**Figure S2. Structure of OspC$_A$ compared to OspC$_B$ and OspC$_K$**. (A) Ribbon diagram showing the homodimeric structure of OspC$_A$ (colored cyan and green) with labelled a-helices and b-strands in the cyan monomer. OspC$_A$ N and C-termini are labelled N and C, respectively. (B) The OspC$_A$ homodimer from the B11-OspC$_A$ complex colored cyan and green superpositioned with OspC$_B$ (PDB ID: 7UJ2) colored dark gray and OspC$_K$ (PDB ID: 7UJ6) colored light gray depicting the structural similarity of each OspC. All OspC's are drawn as Ca-traces.

### Figure S3

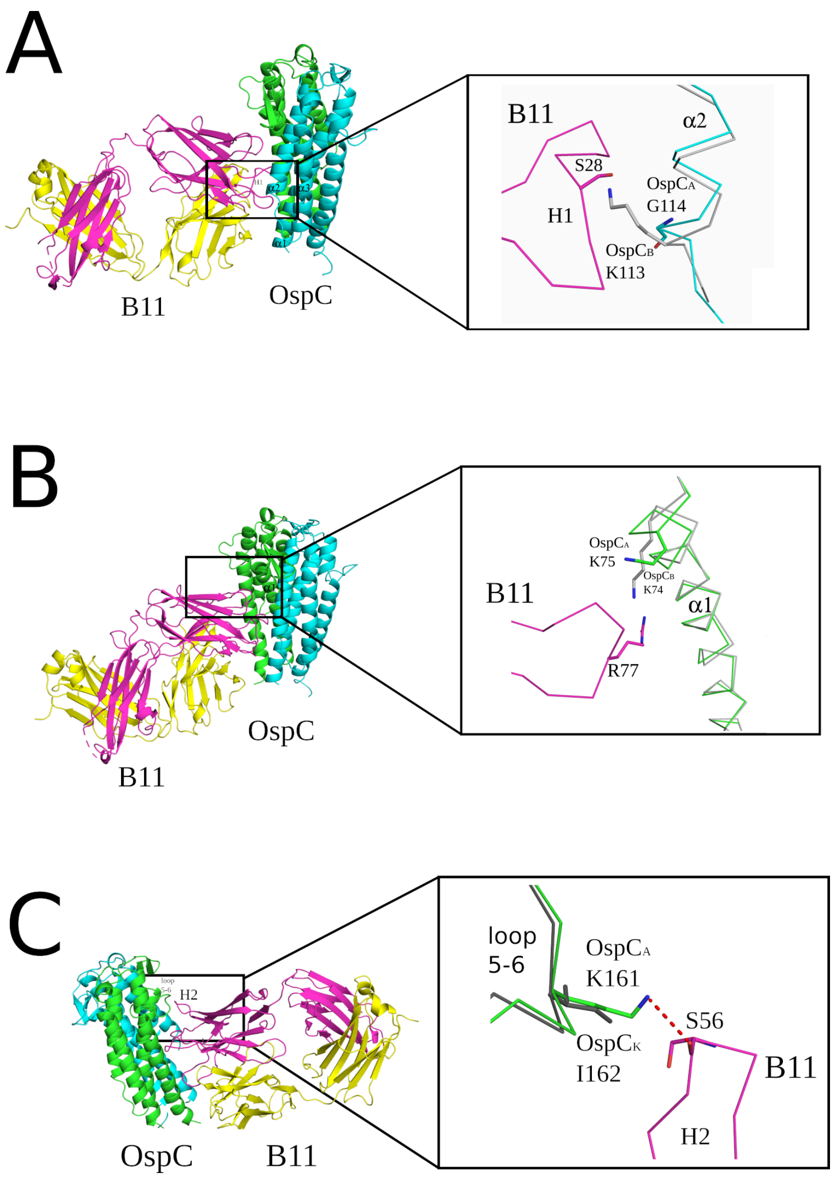


**Figure S3. Structural basis of B11 specificity for $OspC_A$.** Interface between B11 Fab and $OspC_A$ superposed with either $OspC_B$ or $OspC_K$ highlighting the key residues in $OspC_B$ or $Osp_K$ that conceivably diminish interaction with B11. (A) A zoom-out view of the ribbon diagram of the OspC-B11 complex and a zoom-in view of the Ca-traces of $OspC_A$ (cyan) bound to B11 (magenta) with superposed $OspC_B$ (light gray) highlighting the potential steric clash of Lys-113 in $OspC_B$ with Ser-28 in B11. (B) The OspC-B11 ribbon diagram with a closeup of the Ca-traces of $OspC_A$ (green) bound to B11 (magenta) superposed with $OspC_B$ (light gray). The image highlights the proximity of Lys-74 upon superposition of $OspC_B$ revealing the possible electrostatic repulsion with Arg-77 in B11. (C) A zoom-out view of the OspC-B11 complex as a ribbon diagram and a zoom-in perspective of the Ca-traces of $OspC_A$ (green) bound to B5 (magenta) superposed with $OspC_K$ (dark gray). The image shows Ile-162 in $OspC_K$, instead of Lys-161 in $OspC_A$, which precludes H-bonding to B11's Ser-56 as observed in the B11-$OspC_A$ structure. All ribbon diagrams of the OspC-B11 complex are drawn with the $OspC_A$ dimer colored cyan and green. The B11 Fab heavy chain is colored magenta with the light chain colored yellow. Side chains are drawn as sticks and color coordinated to the main chain color, with nitrogen atoms shaded blue and oxygen atoms shaded red. H-bonds are drawn as red dashes.

### Figure S4

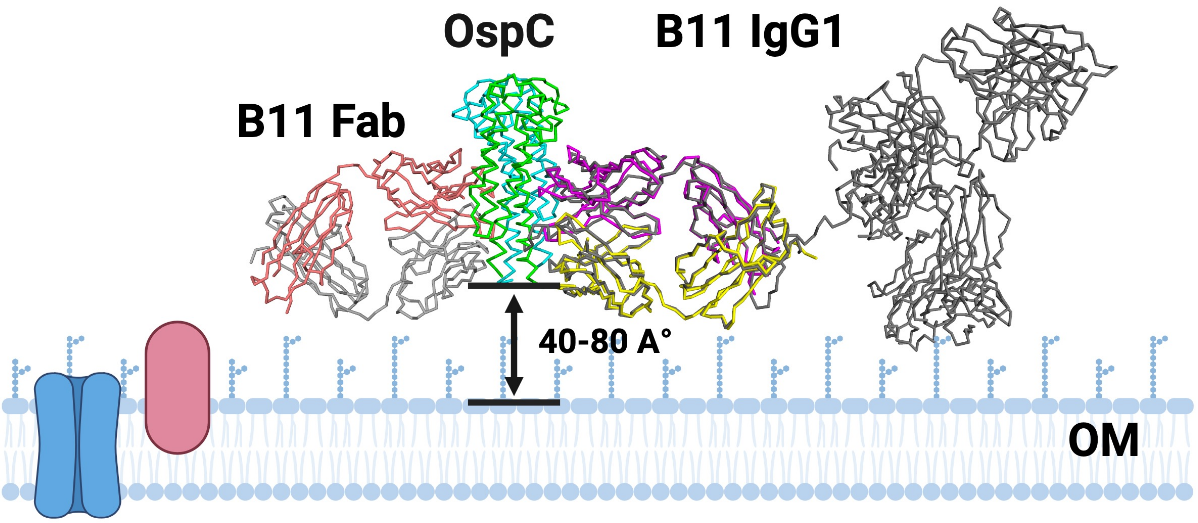


**Figure S4. Proposed alignment of B11 IgG in complex with OspC on the surface of *B. burgdorferi*.** The structure of $OspC_A$ (center; residues 38-201) in complex with B11 Fab (left) and a superpositioned IgG (right) extracted from **Figure 5B** is presented in the context the bacterial outer membrane (OM). OspC is positioned 40-80 Angstroms from the bacterial surface to account for its N-terminal lipid anchor (residues 1-37). In this arrangement, the B11 epitope is accessible to an intact IgG in a side-on orientation. The positioning of OspC at 40-80 Angstroms from the bacterial membrane is based on a model of the *B. turicatae* surface protein 1 (Vsp1) proposed by Lawson and colleagues (J. Bact. 188:4522). Image generated with BioRender, Inc.

### Figure S5

A

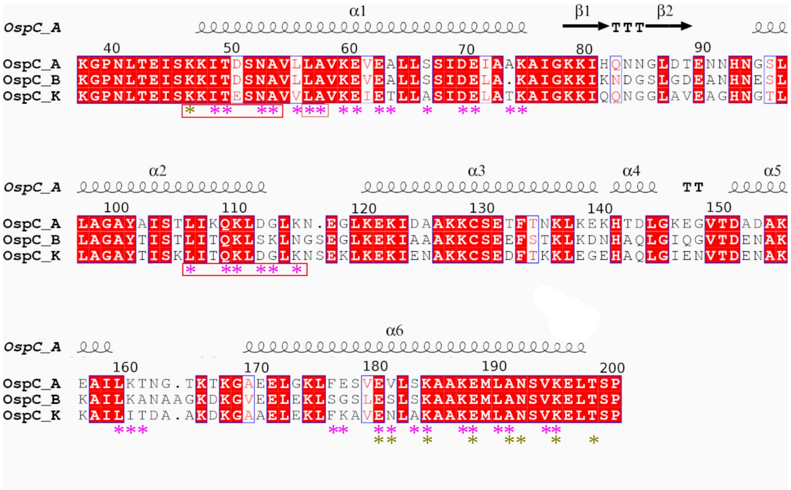


B

```
                                                             *
A     NSADESVKGPNLTEISKKITDSNAVLLAVKEVEALLSSIDEIAAKAIGKKIHQNNGLDTE   90
C3    NSADESVKGPNLAEISKKITESNAVVLAVKEVETLISSIDEIAAKAIGKKIKNDGSLDND   90
I3    NSADESVKGPNLTEISKKITESNAVVLAVKEIETLLSSIDELATKAIGQKIDA-NGLGVQ   89
J     NSADESVKGPNLTEISKKITESNAVVLAVKEIETLLASIDELATKAIGKKIDNNAGLGAE   90
M     NSADESVKGPNLTEISKKITESNAVVLAVKEVETLLASIDEVAKKAIGNLIAQ-NGLNAG   89
      ************:*******:****:*****:*:*::****:* ****: *    .*.

                                   *
A     NNHNGSLLAGAYAISTLIKQKLDGLKN-EGLKEKIDAAKKCSETFTNKLKEKHTDLGKEG   149
C3    ANHNGSLLAGAYAISTLITQKLGGLKNSEELKEKIEEAKKCNKAFTEKLKSSHAELGKQD   150
I3    ADQNGSLLAGAYAISTLITQKLSALNS-ENLKEKVAKVKKCSEDFTNKLKNGNAQLGLAA   148
J     VGQNGSLLAGAYAISTVIIEKLSTLKNVEELKEKITKAKDCSEKFTKKLKDSHAELGKKD   150
M     ANQNGSLLAGAYVISTLIAEKLDGLKNSEELKEKIEDAKKCNKAFTDKLKSSHAELGIAN   149
       .:*********.***:* :**. *:. * ****:  .*.*.: **.***. :::**

                 *            *
A     --VTDADAKEAILKTNGTKTKGAEELGKLFESVEVLSKAAKEMLANSVKELTSPVVAESP   207
C3    --AQDDDAKKAILRTHNTKDKGAEELDKLFKAVENLSKAAKEMLSNSVKELTSPVVAET-   207
I3    --ATDADAKEAILKTNGTKTKGAEELGKLFESVEVLSKAAKEMLANSVKELTSPVVAES-   205
J     --ASDDDAKKAILKTNQANDKGAKELKELFEAVESLSKAAKEMLNKSVKELTSPIVAESP   208
M     GAATDANAKAAILKTNGTKDKGAQELEKLFESVKNLSKAAQETLNNSVKELTSPVVAENP   209
        . * :** ***:*: :: ***:** :**::*: *****:* * :********:***.
```

**Figure S5. OspC sequence alignments.** (A) Structural sequence alignment of OspC types A, B, and K from *B. burgdorferi* depicting the B11-interacting region of $OspC_A$. Olive colored asterisks depict B11 $V_L$ (L1-L3) interacting residues and magenta asterisks depict B11 $V_H$ (H1-H3) interacting residues. Secondary structural elements from $OspC_A$ bound to B11 are illustrated with a-helices 1-6 drawn as black coils and b-strands 1-2 drawn as black arrows above the sequence and labelled accordingly. Red background with white letters connotes sequence identity, white background with red letters connotes sequence similarity. The orange boxes below sequence region 47-54, 56-57, and 107-116 shows B11 contact with the second $OspC_A$ monomer. (**B**) Sequence alignment of OspC types that conceivably bind B11. Primary sequence alignment of the OspC types C3, I3, J, and M with $OspC_A$ underscoring their key sequence similarities with $OspC_A$ that support B11 binding. The red rectangle encapsulates critical residues in a-helix 1 of $OspC_A$ highlighting the region around residue 75 which contains a critical deletion in $OspC_B$ that ostensibly antagonizes B11 binding. Red asterisks above the sequence identify the critical $OspC_A$ residues Gly-114, Lys-161, and Phe-177 which promote interaction with B11 and are sufficiently conserved in OspC types C3, I3, J, and M. Black asterisks below the sequence denote sequence identity with two dots and one dot showing relatively reduced sequence similarity. Figure made with Clusta Omega and ESPript 3.0.
